# The Sea Lamprey Meiotic Map Resolves Ancient Vertebrate Genome Duplications

**DOI:** 10.1101/008953

**Authors:** Jeramiah J. Smith, Melissa C. Keinath

**Affiliations:** Department of Biology, University of Kentucky, Lexington, KY

**Keywords:** Genome, Evolution, Duplication, Vertebrate, Lamprey

## Abstract

Gene and genome duplications serve as an important reservoir of material for the evolution of new biological functions. It is generally accepted that many genes present in vertebrate genomes owe their origin to two whole genome duplications that occurred deep in the ancestry of the vertebrate lineage. However, details regarding the timing and outcome of these duplications are not well resolved. We present high-density meiotic and comparative genomic maps for the sea lamprey, a representative of an ancient lineage that diverged from all other vertebrates approximately 550 million years ago. Linkage analyses yielded a total of 95 linkage groups, similar to the estimated number of germline chromosomes (1N ∼ 99), spanning a total of 5,570.25 cM. Comparative mapping data yield strong support for one ancient whole genome duplication but do not strongly support a hypothetical second event. Rather, these comparative maps reveal several evolutionary independent segmental duplications occurring over the last 600+ million years of chordate evolution. This refined history of vertebrate genome duplication should permit more precise investigations into the evolution of vertebrate gene functions.

## Introduction

It is generally accepted that the ancestral lineage of all or most extant vertebrates experienced two mishaps in cell division that each led to a doubling of the number of chromosomes, and which were inherited by subsequent generations (Kasahara 2007). These duplications are colloquially known as the 2R hypothesis (Holland et al. 1994; Hughes 1999). Although the most common fate of duplicated genes is mutational degradation of one copy (paralog loss), gene duplication also provides raw material for the evolution of new gene functions and the evolutionary tuning of old functions (Ohno 1970; Taylor and Raes 2004; Hahn 2009), thus it seems likely that these duplication events have had a major impact on the early evolutionary trajectory of the vertebrate lineage and establishment of the basal condition from which all vertebrates evolved. Understanding the timing and extent of specific evolutionary events that gave rise to the diverse functionality of vertebrate genes should permit more effective translation of genetic information between invertebrate model species and vertebrate model/target species.

Signatures of ancient vertebrate whole genome duplication (WGD) were apparent in early investigations into the structure and content of vertebrate genomes. Based on genome size, chromosome morphology and isozyme counts, Susumu Onho proposed as early as 1970 that at least one WGD must have occurred prior to the diversification of the ancestral amniote or tetrapod lineage (Ohno 1970). Others have found support for the 2R hypothesis through phylogenetic analysis of gene families and discrete synteny groups that have retained a significant number of genes that presumably derive from the 2R duplications (Larsson et al. 2008; Canestro et al. 2009; Kuraku et al. 2009) but see (Hughes 1999; Asrar et al. 2013), or by examining the counts and distributions of homologous segments within the human genome (Dehal and Boore 2005), among jawed vertebrates (gnathostomes) (Nakatani et al. 2007; Murat et al. 2012) and between gnathostomes and amphioxus (*Branchiostoma floridae*: a cephalochordate) (Putnam et al. 2008). It is important to recognize, however, that analyses of 2R duplication events carry several critical caveats. The 2R duplications are thought to have occurred in the deep ancestry of the vertebrate lineage, in a period that is represented by few extant lineages. Studies comparing tetrapod and fish genomes can reconstruct an ancestral genome that existed ∼400 million years ago (MYA) (Nakatani et al. 2007; Murat et al. 2012) and studies using amphioxus can reconstruct an ancestor that lived ∼600 MYA (Putnam et al. 2008), but it is clear that these respective ancestors substantially pre-and post-date the vertebrate WGD events. As such, analyses of duplication patterns are confounded by fissions, fusions and gene/segmental duplications that have occurred over deep evolutionary time, both prior to and following WGD events.

Lampreys provide a unique perspective on the deep evolutionary history of vertebrate genomes, having diverged from the majority of other vertebrates (i.e. the lineage that gave rise to the gnathostomes) approximately 550 MYA. Comparative studies employing lampreys can therefore be used to resolve the nature and timing of evolutionary events that occurred at or near the base of the vertebrate phylogeny. Analyses of the sea lamprey genome assembly revealed that the duplication content of the lamprey genome is similar to that of other vertebrates and detected patterns of conserved synteny consistent with duplication and extensive paralog loss in both lineages (Smith et al. 2013). Such data are consistent with the most recent duplication event having occurred prior to the divergence of lamprey and gnathostomes, likely predating the evolution of several anciently-derived features such as jaws, paired appendages and neuronal myelination (Smith et al. 2013). Analyses of a second lamprey genome (*Lethenteron japonicum)* provided further confirmation that duplication content of lamprey genomes is similar to that of jawed vertebrates and suggested the possibility that the lamprey lineage had undergone a third whole genome duplication, subsequent to 2R (discussed in more detail below) (Mehta et al. 2013). However, it is important to recognize that both of these studies relied on the conventional wisdom that gnathostomes have undergone two rounds of WGD and did not explicitly test alternate evolutionary models. This is largely due to the fact that existing lamprey assemblies do not currently permit the resolution of chromosome-scale patterns of homology or the robust reconstruction of ancestral vertebrate chromosomes, which are necessary to test several key aspects of the 2R hypothesis.

To better resolve the chromosome-scale structure of the lamprey genome, we constructed the first meiotic map for a lamprey species by restriction site associated DNA sequencing (RADseq) (Miller et al. 2007; Baird et al. 2008; Amores et al. 2011) of siblings from a controlled cross between two wild-captured *Petromyzon marinus*. Sequence information from this mapping family permitted the anchoring of approximately half of the lamprey genome assembly to the larger chromosomal structure of the genome. Analysis of chromosome-scale patterns of conserved synteny between lamprey and gnathostomes permits the identification of an ancestral complement of chromosomes in the pre-duplication lineage and reveals the effects of duplication in both lineages. Reconstructed ancestral chromosomes largely recapitulate a complement of chromosomes that has been previously proposed for the ancestral vertebrate lineage prior to the first WGD (1R) (Nakatani et al. 2007; Murat et al. 2012) and include three additional homology groups that likely represent additional ancestral chromosomes. While the patterns of conserved synteny in lamprey/gnathostome comparative maps strongly support the existence of the 1R duplication, they lend less support to a hypothetical second duplication (2R), revealing ancient fissions and fusions that have dispersed the signal of 1R in extant vertebrate genomes and specific segmental duplications that gave rise to classically described 2R-derived synteny regions.

## Results

### Linkage mapping and analysis

Our RADseq approach yielded 7,215 segregating markers that could be directly assigned to a scaffold from the lamprey genome assembly (AEFG01 : http://www.ncbi.nlm.nih.gov) or further aided in the merger of maternal and paternal maps. The ability to characterize large numbers of markers of diverse segregation phase was facilitated in part by a relatively high frequency of segregating polymorphism, characteristic of *P. marinus* (Smith et al. 2013), and permitted the construction of both parent-specific and parent-averaged linkage maps. In total, 5,275 markers could be confidently placed within linkage groups containing at least 10 markers linked at a minimum LOD score (log of odds) of 3.0, yielding a total of 95 linkage groups (LGs), which is similar to the estimated number of germline chromosomes (1N ∼ 99) (Smith et al. 2010). These linkage groups span a total of 5,570.25 cM of the parent-averaged map at an average marker density of one marker per 1.3 cM (Figures S1 & S2, Table S1). In total 50.3% of orthology informative genes present in the somatic genome assembly (5483 of 10891 in the lamprey/chicken comparative map) and 38.6% of all non-gap sequence (including 94 of the 100 longest scaffolds) could be placed on the 95 primary linkage groups. The high marker density and cross validation of maternal and paternal genetic maps provided by this approach yielded robust chromosome-scale scaffolding of the lamprey genome assembly and comparative maps (Tables S2 and S3).

### Genome-wide patterns of conserved synteny

Analysis of lamprey/gnathostome (human and chicken) comparative maps reveals statistically significant enrichment of presumptive orthologs on several pairs of lamprey/gnathostome chromosomes (Figure S3). Such patterns are indicative of relatively strong conservation of chromosome-scale synteny across deep evolutionary time. Notably, a majority of the lamprey LGs that showed a significant enrichment of homologs from one gnathostome (chicken) chromosome also showed a significant enrichment of homologs from a second chicken chromosome (61%). This pattern was even more prominent among the 30 largest linkage groups, with 90% of these linkage groups showing statistically significant enrichment of orthologs from two or more chicken chromosomes. Likewise, 93% of chicken chromosomes that showed a significant enrichment of homologs from a lamprey LG showed a significant enrichment of homologs from a second lamprey LG. Similar patterns are also visible in the lamprey/human comparative map, albeit obscured by the effects of increased rates of intrachromosomal rearrangement in the mammalian lineage (ICGSC 2004; Smith and Voss 2006; Alfoldi et al. 2011; Voss et al. 2011; Amemiya et al. 2013) (Figure S3). Given the assumption that selective convergence in chromosomal gene content is not a major driving force in vertebrate karyotype evolution, enrichment of homologs between chicken and lamprey is interpreted as strong evidence that the respective gnathostome and lamprey chromosomes/chromosomal segments are derived from a single common ancestral chromosome.

The ancestral groupings that are resolved by the lamprey/gnathostome comparative maps largely recapitulate a set of ten ancestral chromosomes previously proposed for the pre-1R ancestor (ancA-J: Table 1) (Nakatani et al. 2007; Murat et al. 2012). In addition to these ten previously identified chromosomes, the lamprey/chicken comparative map reveals three additional clusters of orthology that likely represent independent chromosomes from the presumptive pre-1R ancestor, suggesting an ancestral chromosome number of ∼13 (Figure 1). Regions of non-independence between presumptive ancestral orthology groups (i.e. overlaps of highlighted regions on the x-or y-axes of Figure 1) reflect fusion or translocation events that occurred in the 550 million years since the divergence of lamprey and gnathostome lineages or false mergers of linkage groups/scaffolds. The effect of these factors is relatively minor with respect to the lamprey/chicken comparative map, suggesting relatively low rates of interchromosomal rearrangement in both lineages and fidelity of ortholog identification. Barring other patterns within the data, the observed distribution of ancestrally syntenic regions across multiple derived chromosomes could result from fissions and translocations or from large-scale duplication. However, it is possible to more precisely reconstruct the evolutionary history of derived chromosomes by examining the distribution of orthologous loci among ancestrally associated segments.

**Table 1:** Correspondence between ancestral (pre-1R) chromosomes identified in this study and others.

| Current Study | Nakatani *et al*<sup>11</sup> | Putnam *et al*<sup>13</sup> |
| --- | --- | --- |
| A | A | 1,2 |
| B | B | 3,4 |
| C | C | 6,7 |
| D | D | 13 |
| E | E | 16 |
| F | F | 8,9,10 |
| G | G | 11,14 |
| H | H | 17 |
| I | I | 15 |
| J | J | 5 |
| K | F | - |
| L | A | 9 |
| M | - | - |

**Figure 1.**
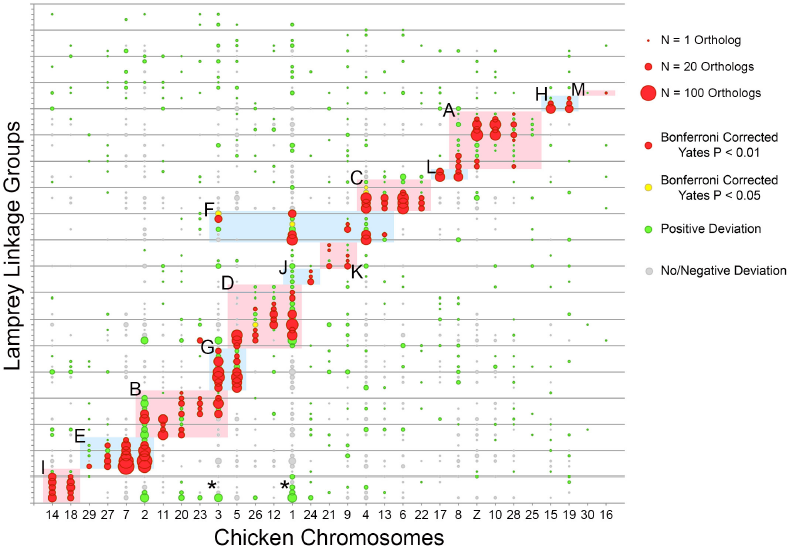
Distribution of conserved syntenic genes in lamprey and chicken genomes. The size of each circle is proportional to the number of orthologous genes located on the corresponding lamprey LG and chicken chromosome. The color of each circle represents the degree to which the number of observed orthologs deviates from null expectations under a uniform distribution across an identical number of LGs, chromosomes and genes per LG and chromosome. Shaded regions of the plot designate homology groups that correspond to presumptive ancestral chromosomes, marked A - M (Extended Data Table 1). An example of an ancient pre-1R duplication involving a subsegment of ancI is marked with asterisks. The ordering of lamprey LGs along the y-axis is provided in Table S4.

In both gnathostomes and lamprey, the derivatives of pre-1R chromosomes are typically distributed across two or more chromosomes, with the exception of one small orthology group that is represented by a single lamprey LG (“M” in Figure 1). With respect to the chicken genome, a majority of orthology groups show a 1:2 ratio of ancestral:derived (a:d) chromosomes (N = 7 for ancG - M). Three other orthology groups (ancA, C and E) show ∼1:4 ratios, consisting of two macrochromosomes and two microchromosomes. The three remaining orthology groups (ancB, D and F) show more complex patterns consistent with fissions and small duplications having occurred subsequent to 1R, the derivatives of which show clear breaks in synteny suggesting that these groups formerly constituted 1a:2d or 1a:3d ratios (Figure S4). We interpret the presence of an overarching 1:2 (or more) ratio of ancestral to derived gnathostome chromosomes as consistent with a scenario wherein all 13 ancestral chromosomes were affected by at least one WGD.

### Genomic distribution of orthologs and paralogs

To further evaluate the patterns of duplication that are revealed by lamprey comparative maps, we examined the distribution of orthologies within lamprey and chicken derivatives of the 13 ancestral chromosomes. The extensive loss and (rarer) retention of paralogs yield distinctive genomic signatures in the conserved syntenic structure of duplicated chromosomes. The independent nature of paralog loss results in a situation wherein two derived chromosomes retain a unique subset of genes, which form an interdigitated pattern when mapped to their homologous (unduplicated) chromosome (Amores et al. 2011; Smith et al. 2013). Lamprey/gnathostome comparative maps show a similar, but somewhat more complex pattern of conserved synteny. Specifically, gnathostome (chicken) chromosomes retain unique subsets of genes located on several lamprey chromosomes and lamprey chromosomes retain unique subsets of genes located on several (typically two) gnathostome chromosomes (Figure 2). We interpret such a pattern as consistent with divergence of basal lamprey and gnathostome lineages shortly after WGD, such that rediploidization occurred through largely independent series of mutational events in the two lineages.

**Figure 2.**
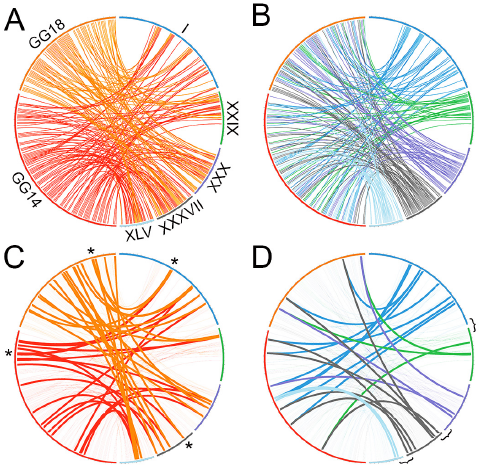
Distribution of individual orthologous genes across paralogous chicken and lamprey chromosomes. A) Orthologs from chicken chromosomes GG14 and GG18 are distributed across five lamprey LGs (the complete set of lamprey linkage groups with significant enrichment for orthologs on these two chicken chromosomes). B) Orthologs from these LGs are distributed across GG14 and GG18 (the complete set of chicken chromosomes with significant enrichment for orthologs on these lamprey linkage groups). C) Orthologies plotted in A, with retained chicken paralogs denoted by bold lines. D) Orthologies plotted in panel B, with retained lamprey paralogs denoted by bold lines. Roman numerals designate lamprey LGs. Asterisks mark the location of four conserved paralogs that are derived from a single gene located on ancI. Brackets denote presumptive ancestral linkages that are supported by the distribution of paralogous genes.

The distribution of retained paralogs also supports a scenario of shared duplication and independent divergence. Broad-scale information provided by the lamprey linkage map allows us to directly trace the ancestry of these paralogs to their pre-duplication chromosome, thereby confirming their evolutionary diversification via large-scale (whole genome) duplication. We detect synteny-supported paralogs on lamprey and chicken derivatives of all proposed ancestral chromosomes, with the exception of the apparently gene-poor ancM. Given that ∼1/2 of the lamprey genome is anchored to the linkage map, counts of retained paralogs in lamprey (N=266) and chicken (N=503) indicate that the derivatives of the 13 ancestral chromosomes have retained similar numbers of duplicated genes, consistent with previous studies (Smith et al. 2013). The distribution of retained paralogs in lamprey and chicken genomes can be further leveraged to resolve fission events that have fractured chromosomes subsequent to an ancestral WGD (Figure 2D); such fission events appear to have contributed significantly to the karyotypic evolution in the ancestral lineage that gave rise to the northern hemisphere lampreys (including *P. marinus*) (Hardisty 1971).

### Hypothesis testing

The identification of ancestral chromosomes and their derivative genomic segments provides a framework for testing hypotheses as to the origin of duplicated segments in gnathostome genomes. Below we address three previously proposed explanations for the distribution of paralogous regions in gnathostome genomes: **A)** two rounds of WGD (either before or after the divergence of the lamprey lineage) with essentially random loss of duplicated genes (Ohno 1970; Kasahara 2007), **B)** two rounds of WGD followed by extensive loss of large duplicated segments/chromosomes (Nakatani et al. 2007; Murat et al. 2012), **C)** only segmental duplication (Asrar et al. 2013) and one WGD with additional segmental duplications pre-or post-dating WGD.

Observed frequencies of a:d chromosomes deviate strongly from expectations under the strict 2R model, even given a liberal interpretation that all 1a:>2d ratios (N=5) are supportive of 2R (Table 2, G_ajd(c)_=35.199, df=1, p=2.98E^-9^). However, large-scale chromosome loss has been invoked in some reconstructions as a significant contributing factor to post-2R assortment of chromosomal segments (Nakatani et al. 2007; Murat et al. 2012). We reasoned that the plausibility of the large-scale deletion model (and other models) might be tested by measuring the degree to which the distribution of losses (or gains in later tests) across ancestral orthology groups conforms to expected frequencies under a random mutational model (i.e. distributed Poisson for rare events). Depending on the timing of loss relative to 1R and 2R, the number of losses that are necessary to generate the observed patterns ranges from 9, when all 1a:2d ratios are considered to be the product of the product of deletion of a 1R paralog (“conservative” model), to 18 when 1a:2d ratios are considered to be the product of the product of recurrent deletions of different paralogous segments post 2R (“liberal” model, Table 3). It is worth noting that in this context “loss” may also include large-scale degradation or more generally, failure to identify derivative segments.

**Table 2:** Comparison of observed and expected patterns of duplication under 2R (two whole genome duplications).

| # Dup. | Obs. | Exp. | Exp. (adj.) | G | P-value |
| --- | --- | --- | --- | --- | --- |
| 1 | 8 | 0 | 0.5 | 35.20 | 2.98E<sup>-9</sup> |
| >1 | 5 | 13 | 12.5 |  |  |

Dup – Duplications, Obs. – observed, Exp. – expected, adj. – expected values after continuity correction of expected values in order to permit calculation of conservative G-tests

**Table 3:** Observed numbers of duplicated or deleted segments under various models of chromosomal evolution.

| Ancestral chromosome | Conservative count of deletions | Liberal count of deletions | Conservative count of duplications | Liberal count of duplications |
| --- | --- | --- | --- | --- |
| A | 0 | 0 | 2 (1) | 2 (1) |
| B | 0 | 1 | 2 (1) | 3 (2) |
| C | 0 | 0 | 2 (1) | 3 (2) |
| D | 1 | 1 | 2 (1) | 3 (2) |
| E | 0 | 0 | 2 (1) | 2 (1) |
| F | 1 | 2 | 1 | 1 |
| G | 1 | 2 | 1 | 1 |
| H | 1 | 2 | 1 | 1 |
| I | 1 | 2 | 1 | 1 |
| J | 1 | 2 | 1 | 1 |
| K | 1 | 2 | 1 | 1 |
| L | 1 | 2 | 1 | 1 |
| M | 1 | 2 | 1 | 1 |
| Total | 9 | 18 | 18 (5) | 21 (8) |

See text for further details regarding the annotation of duplication events. Numbers in parentheses reflect counts in excess of those accounted for by one whole genome duplication.

Corresponding statistical tests for goodness of fit reveal that the distribution of deletion events across ancestral chromosomes is inconsistent with expected patterns, under both the strict liberal and conservative mutational models (Table 4; conservative: G_ajd(W)_=7.819, df=2, p=0.020; liberal: G_ajd(W)_=7.819, df=2, p=0.006). However, by considering all possible permutations of liberal and conservative counting schemes across the set of chromosomes with 1a:2d ratios, it is possible to identify a range of scenarios wherein observed counts do not reject a random mutational model. The best fit to the “2R plus deletion” model requires 12 chromosomal losses, with three recurrent losses of paralogous chromosomes post-2R (Table 4; optimal permutation: G_ajd(W)_=2.349, df=2, p=0.309). The simplest model failing to reject an underlying Poisson distribution at p<0.05 involved 10 chromosomal losses, with one recurrent loss.

**Table 4:** Comparison of observed and expected patterns of duplication deriving from two rounds of whole genome duplication and large-scale segmental or chromosomal loss

| # Del | Conservative Model |  |  | Liberal Model |  |  | Optimal Permutation |  |  |
| --- | --- | --- | --- | --- | --- | --- | --- | --- | --- |
|  | Obs | Exp | G / P-value | Obs | Exp | G / P-value | Obs | Exp | G / P-value |
| 0 | 4 | 6.51 | 7.82 | 3 | 3.26 | 10.33 | 4 | 5.16 | 2.35 |
| 1 | 9 | 4.50 | 2.01E-02 | 2 | 4.51 | 5.72E-03 | 6 | 4.77 | 0.31 |
| 2 | 0 | 1.56 |  | 8 | 3.12 |  | 3 | 2.20 |  |
| >2 | 0 | 0.43 |  | 0 | 2.12 |  | 0 | 0.87 |  |

Counts of observed duplicates under conservative and liberal models are provided in Table 3. Dup - Duplications, Obs. - observed, Exp. - expected.

A similar scheme can be used to assess the distribution of duplications under a scenario wherein observed duplications are strictly the product of sporadic duplication events that individually affect only single chromosomes or large chromosomal segments. As above, the numbers of duplication events necessary to recover patterns in the lamprey/gnathostome comparative map were estimated using conservative (down-weighting recurrent events) and liberal (up-weighting recurrent events) schemes. Under the “segmental duplication only” model, all counting schemes and permutations thereof yield distributions that reject a random mutational model (Tables 3 & 5; conservative set: G_ajd(W)_=12.601, df=3, p=0.006; liberal set: G_ajd(W)_=10.274, df=3, p=0.016; best fit permutation: G_ajd(W)_=9.415, df=3, p=0.024). Thus segmental/chromosomal duplication appears to be insufficient, in itself, to explain observed patterns of duplication.

**Table 5:** Comparison of observed and expected patterns of duplication deriving from segmental and chromosomal duplication.

| # Dup | Conservative Model |  |  | Liberal Model |  |  | Optimal Permutation |  |  |
| --- | --- | --- | --- | --- | --- | --- | --- | --- | --- |
|  | Obs | Exp | G / P-value | Obs | Exp | G / P-value | Obs | Exp | G / P-value |
| 0 | 0 | 3.26 | 12.6 | 0 | 2.58 | 10.27 | 0 | 3.01 | 9.42 |
| 1 | 8 | 4.51 | 5.58E-03 | 8 | 4.18 | 1.64E-02 | 8 | 4.41 | 2.43E-02 |
| 2 | 5 | 3.12 |  | 2 | 3.37 |  | 4 | 3.22 |  |
| 3 | 0 | 1.44 |  | 3 | 1.82 |  | 1 | 1.57 |  |
| >3 | 0 | 0.68 |  | 0 | 1.05 |  | 0 | 0.79 |  |

Counts of observed duplicates under conservative and liberal models are provided in Table 3. Dup - Duplications, Obs. - observed, Exp. - expected.

Finally, this same logical framework was also used to test the fit of distributions deriving from a single WGD event, with segmental duplications either preceding or following the event. After accounting for 1R, the distribution of remaining presumptive segmental duplicates is consistent with (i.e. fails to reject for all counts and permutations) a model of sporadic segmental/chromosomal duplication (Tables 3 & 6; conservative: G_ajd(W)_=2.036, df=2, p=0.361; liberal: G_ajd(W)_=3.534, df=2, p=0.171; optimal permutation with one recurrent event: G_ajd(W)_=0.308, df=2, p=0.857), with permutations with duplication pre-and post-dating a single WGD showing the smallest deviation from expectations under a random mutation model.

**Table 6:** Comparison of observed and expected patterns of duplication deriving from segmental and chromosomal duplication, in addition to one round of whole genome duplication.

| # Dup | Conservative Model |  |  | Liberal Model |  |  | Optimal Permutation |  |  |
| --- | --- | --- | --- | --- | --- | --- | --- | --- | --- |
|  | Obs | Exp | G / P-value | Obs | Exp | G / P-value | Obs | Exp | G / P-value |
| 0 | 8 | 8.85 | 2.04 | 8 | 7.03 | 3.53 | 8 | 8.19 | 0.31 |
| 1 | 5 | 3.40 | 0.36 | 2 | 4.32 | 0.17 | 4 | 3.78 | 0.86 |
| 2 | 0 | 0.65 |  | 3 | 1.33 |  | 1 | 0.87 |  |
| >2 | 0 | 0.09 |  | 0 | 0.32 |  | 0 | 0.15 |  |

Counts of observed duplicates under conservative and liberal models are provided in Table 3. Dup - Duplications, Obs. - observed, Exp. - expected.

In summary, two classes of models failed to reject expectations under a model of random mutation: 1) those invoking two rounds of whole genome duplication and between 10 and 16 independent chromosomal losses and 2) those invoking a single round of whole genome duplication and 5 - 6 segmental/chromosomal duplications.

## Discussion

Taken together, analyses of lamprey/gnathostome comparative maps resolve and refine the complement of chromosomes that existed in the pre-1R vertebrate ancestor and indicate that the divergence of the two basal vertebrate lineages (that respectively gave rise to lampreys and gnathostomes) occurred shortly after a single whole genome duplication event. Although comparative maps do reveal a clear global signature of one whole genome duplication event, we do not observe global signatures consistent with a second WGD in the ancestry of the gnathostome lineage. Rather, these comparative maps appear to reveal alternate evolutionary origins of paralogous regions that have been previously cited in support the 2R duplication. Specifically, these patterns of conserved synteny are consistent with previously cited examples of fourfold conserved synteny (including HOX paralogy regions) being the product of ancient segmental duplication, followed by a single round of WGD.

Although the evolutionary model inspired by analysis of the lamprey linkage map differs from more classical versions of the 2R model that have been supported by other studies, the “1R plus segmental duplication” model does not seem to be at odds with the primary findings of those studies. For example, studies examining the distribution and depth of syntenic paralogous regions in the human genome estimated that each region of the human genome contained paralogous copies the existed in at least three other distinct locations (Dehal and Boore 2005; Putnam et al. 2008). Our analyses reveal that extensive intrachromosomal rearrangement has occurred post-1R (Figure 1S4) and corroborate numerous studies demonstrating that rates of interchromosomal rearrangement are much higher in mammals than in other vertebrate lineages (Figure 1S3) (ICGSC 2004; Smith and Voss 2006; Alfoldi et al. 2011; Voss et al. 2011; Amemiya et al. 2013). As discussed above, duplication and fission/translocation events can both distribute orthologous segments in similar patterns, and the same holds true for paralogous segments within a genome. It seems clear that interchromosomal rearrangement has played a major role in structuring the distribution of conserved syntenic regions between pre-1R ancestral chromosomes and their human derivatives, as can be seen by comparing Figures 1 and 1S3. Thus, patterns of conserved synteny in the human genome do not seem to be at odds with the findings of the current study.

Other studies have leveraged comparisons among gnathostomes (Nakatani et al. 2007; Murat et al. 2012) or between gnathostomes and chordates (Putnam et al. 2008) to reconstruct pre-1R orthology groups. Notably, pre-1R chromosomes reconstructed via the lamprey meiotic map closely resemble those proposed by these previous studies. However, these studies did not explicitly address alternate evolutionary models or departures from 1a:4d (expected under 2R). Still other studies have focused on a smaller number of genomic regions that are thought to represent the largest and best-conserved regions of 4-fold paralogy (Larsson et al. 2008; Canestro et al. 2009; Kuraku et al. 2009), although the existence of these well-defined regions is consistent with several proposed models. Finally, two recent studies (including a study that was co-authored by one of us) used draft lamprey genomes to examine the relative timing of WGD and divergence events, but relied on the assumption that gnathostome genomes were indeed the product of two rounds of whole genome duplication (Mehta et al. 2013; Smith et al. 2013). In general, the “1R plus segmental duplication” model appears to be consistent with primary findings of all studies of 2R performed to date (discussed in detail below), though it seems that this alternate model was not readily apparent in the absence of chromosome-scale data from lamprey.

### Chromosome loss vs. segmental duplication

As described above, we interpret patterns that are visually apparent in the lamprey/chicken comparative map as consistent with being generated by one WGD with additional large paralogy regions being the product of rare segmental duplications occurring both before and after WGD. Moreover, the distribution of duplicated segments under this scenario is consistent with expectations given a simple random mutational model. Other scenarios invoking 2 rounds of WGD followed by chromosomal loss also yield counts that are consistent with a random mutational model but require a substantially larger number of mutational steps to conform to the model. Specifically, models invoking one WGD require as few as six mutational steps (one WGD plus five segmental duplications/fissions), whereas models invoking two WGD events require between 12 and 18 steps (14 for the optimal permutation: two WGD events and 12 deletions). A scenario of one WGD pre-and post-dated by sporadic segmental duplications appears to provide the most parsimonious explanation for the distribution of large paralogous segments across gnathostome genomes. Below we discuss some of the salient evolutionary details of this scenario and relate these to previous observations that were conceptualized under the 2R hypothesis.

Our analyses indicate that both WGD and large segmental duplications likely played key roles in the deep history of vertebrate genome evolution. Among the duplication and fission events apparent in the comparative map, one recurrent pattern is particularly striking. All of the statistically supported groupings of ancestrally-associated chicken chromosomes that deviate from a pattern of 1a:2d chromosomes share a common feature in that they involve groups of macro-and micro-chromosomes. Previous studies have shown that microchromosomes are present in most major gnathostome lineages and that they represent distinct evolutionarily conserved entities at least to the base of the tetrapod lineage (∼350 MYA) (Voss et al. 2011; Uno et al. 2012). Several of these microchromosomes: GG28 (ancA), GG20, 23 (ancB), GG13, 22 (ancC), GG26 (ancD) and GG27, 29 (ancE), bear the signature of duplication and fission events that are temporally distinct from 1R. The “single WGD” model identifies three sets of macro-and micro-chromosomes that experienced duplication prior to 1R (specifically, derivatives of ancA, C, and E). Notably, these three proposed duplications account for essentially all of the four-fold paralogous conserved syntenies that have been classically studied in the context of the 2R hypothesis, including large synteny groups that are exemplified by paralogs of HOX and RAR (ancE), MHC and ALDH1 (ancA) and NPYR (ancC) loci (Larsson et al. 2008; Canestro et al. 2009; Kuraku et al. 2009). Previous reconstructions of the ALDH1-syntenic region also strongly implicate a pre-1R intrachromosomal duplication of ancA followed by chromosomal fission (Canestro et al. 2009), consistent with the single WGD model.

Patterns of conserved synteny in ancC and ancE suggest plausible mechanisms underlying the pre-1R duplication of these 1a:4d orthology groups that are distinct from those previously described for ancA (Canestro et al. 2009). The observed pattern for ancE is consistent with one of the expected segregation products following a Robertsonian fusion between ancestral Hox-bearing chromosome and another (presumably larger) chromosome. Following such a fusion, normal disjunctions can give rise to two chromosomes that are respectively similar to the ancE-derived micro- (GG27, 29) and macro-(GG2, 7) chromosomes, with the larger possessing material duplicated from the smaller. A similar mechanism seems equally viable for the ancE (NPYR-bearing) chromosomes, although other mutations could yield similar patterns of duplication (Moore and Best 2001). The proposed scenario of duplication of the ancestral Hox-bearing chromosome early in the course of vertebrate evolution (before 1R) is consistent with the presence of four Hox clusters in gnathostomes and recent studies either supporting (Smith et al. 2013) or confirming (Mehta et al. 2013) the existence of more than two paralogous Hox clusters in lamprey genomes. The segmental duplication model is therefore consistent with both well-defined mutational mechanisms and studies confirming the existence of paralogous conserved syntenies shared by gnathostome and lamprey lineages. Altogether, we interpret patterns of conserved synteny between lamprey and gnathostomes as strongly supporting the occurrence of a single WGD predating the lamprey/gnathostome split, with other paralogous regions (in excess of 1a:2d) being the product of a small number of independent segmental duplications.

The above interpretation provides a simple explanation that integrates observations from lamprey/gnathostome comparative maps with previous analyses of vertebrate syntenic regions. However, we acknowledge that the simplest explanation may not always be correct. If microchromosome-associated duplications are, in reality, the product of a gnathostome-specific WGD (i.e. 2R), then the evolutionary assortment of post-2R chromosomes must have progressed in a manner that is substantially different from other vertebrate whole genome duplication events [i.e. in the *Xenopus* lineage (Evans et al. 2004; Uno et al. 2013) in the salmonid lineage (Berthelot et al. 2014) or near the base of the teleost fish lineage (Amores et al. 2011)]. Under a hypothetical 2R, the evolutionary assortment of post-2R chromosomes would have involved large-scale loss of chromosomes and chromosomal segments, rather than the gradual diversification and nearly random mutational loss of paralogs. Analyses of other vertebrate lineages that have undergone more recent large-scale duplication and chromosomal loss might ultimately reveal a candidate mechanism for post-2R loss.

It is also notable that our analyses treat chromosome losses and segmental duplications as functionally equivalent with respect to their mutational effect. It is important to recognize, however, that the fitness effects of a specific mutation (whole genome duplication, deletion, segmental duplication or otherwise) are contingent on the genetic background and environment in which the mutation occurred, with the probability of fixation being further contingent on the population structure. Studies examining the effects of segmental duplication and chromosome loss across diverse vertebrate taxa will ultimately provide some insight into the relative probabilities of fixation, but it seems certain that these parameters will never be known for the ancestral vertebrate lineage. Given the relative simplicity of incorporating previous observations into the model, we assert that the distribution of paralogous segments in vertebrate genomes is currently best explained by one WGD and the evolution of microchromosome-associated paralogy regions via segmental duplication.

## Ancestral vs. Independent Duplication

Two recent studies (Mehta et al. 2013; Smith et al. 2013) provide strong evidence that ancient WGD(s) impacted the lamprey genome, and that subsequent paralog losses progressed largely independently in lamprey and gnathostome ancestors. However, these studies differ in their interpretation of duplication patterns relative to the divergence of the basal lineages that gave rise to lampreys and gnathostomes. It should be noted that the timing of these ancient duplication and divergence events does not strongly affect the identification of ancestral chromosomes in the current study. The observation of non-independent distributions of paralog losses was considered evidence that duplication preceded divergence (Smith et al. 2013), whereas differences in the 4DTv (transversions at fourfold degenerate sites) distributions for lamprey vs. gnathostome paralogs were considered evidence for independent duplication in the lamprey lineage (Mehta et al. 2013). The current study verifies that estimates of paralog retention/loss rates in anc-derived paralogous regions are consistent with those estimated from the entire genome assembly, lending some support to the idea that divergence followed duplication. Moreover, it should be noted that estimates of 4DTv are highly contingent on patterns of substitution (Tang et al. 2008) and that long-term substitution bias in the lamprey lineage is known to have driven the lamprey genomes to exceedingly high GC content especially within coding regions (Qiu et al. 2011; Mehta et al. 2013; Smith et al. 2013). This bias (although not the GC content itself) is seemingly sufficient to explain variation in 4DTv between lampreys and gnathostomes. Indeed, greater degrees of intraspecific variation in 4DTv have been observed among dicot plants in the wake of the ancestral γ hexaploidization event (Tang et al. 2008). We interpret the bulk of available evidence as indicating that 1R (and a finite number of large segmental duplications) predated the divergence of the basal lineages that gave rise to lampreys and gnathostomes, with a relatively small number of large segmental duplications occurring subsequent to 1R. It seems possible, however, that material deleted from somatic cells during programmed genome rearrangement (Smith et al. 2009; Smith et al. 2012) (and therefore not represented in the current study), could be the product of large-scale duplication (or more limited duplication) in the ancestral lamprey/cyclostome lineage. We anticipate that the development of genome assemblies for other taxa (especially germline genomes from hagfish and divergent lamprey species) will improve the temporal resolution of ancestral duplication and divergence events.

By resolving the deep evolutionary history of vertebrate genomes, analyses leveraging the anchored lamprey genome provide new context for understanding the ancestry and evolutionary diversification of vertebrates. We anticipate that incorporation of a more historically accurate duplication history into future analyses, and revision of previous evolutionary studies that have conceptualized vertebrate evolution in light of other hypothesis (including 2R), will provide a more robust understanding of the functional evolution of vertebrate genomes and aid in translating information between vertebrates and non-vertebrate species.

## Materials and Methods

### RadSeq Genotyping

Lamprey embryos were produced by in vitro fertilization (Nikitina et al. 2009) of eggs from a single female with sperm from a single male. DNA was extracted from 141 lamprey embryos (21 days post fertilization) via standard phenol/chloroform extraction (Sambrook and Russell 2001) and evaluated by gel electrophoresis. A total of 30 samples, containing highly intact DNA were selected for RADseq, along with parental DNAs that were extracted from muscle tissue. RADseq was performed by Floragenex, Inc. (Eugene, Oregon), yielding a total of 147 million 95bp sequence fragments (3,590,251 – 5,489,313 per individual) anchored to SbfI restriction sites. The *Stacks* software package (Catchen et al. 2011) was used to identify segregating polymorphisms and generate genotype calls for parents and offspring. For this study, the average depth of coverage per locus was 187.3 reads (range 156.0 – 217.3 per individual). Because sequence information is generated from DNA extracted from whole embryos, we expect to primarily sample DNA from somatic cells. Therefore the current dataset is missing approximately 20% of the germline genome and 1-2000 genes that are specifically retained in the genome of lamprey germ cells (Smith et al. 2009; Smith et al. 2012). Similarly, the method may also fail to produce genotypes from genomic regions with high insertion/deletion frequencies. Given the relatively small number of genes that are eliminated by programmed genome rearrangement, we do not expect exclusion of these regions to systematically bias our comparative analyses especially as they relate to the distribution of duplicated segments in gnathostome genomes.

### Linkage Analysis

Linkage analysis was performed via maximum likelihood mapping using JoinMap software package and default parameters, except that the number of optimization rounds was increased from three to five to ensure accurate internal ordering of large numbers of tightly linked markers (Stam 1993; Van Ooijen 2011). In order to circumvent limitations of the software package related to computational overhead, efficiently anchor the map to the existing assembly and permit robust integration of female and male maps, we limited our analyses to markers that, 1) aligned to the published genome assembly [>95% identity over ≥90 bp via *BLAST* (Altschul et al. 1990)], 2) yielded informative segregation phasing and 3) yielded genotypic information for at least 20 of 30 offspring. These were further supplemented with markers that were specifically informative for one or both parents, regardless of alignment to the genome assembly, permitted that they were genotyped for at least 27 of 30 offspring for biallelic markers [llxlm, nnxnp or hkxhk (Maliepaard et al. 1997; Van Ooijen 2011)] or at least 25 of 30 offspring for tri-and tetra-allelic markers [efxeg or abxcd (Maliepaard et al. 1997; Van Ooijen 2011)]. The maximum likelihood mapping algorithm for mapping of a full sib family from outbred parents (CP) (Van Ooijen 2011), was used to generate a linkage map from a total of 7,215 potentially informative markers. Linkage groups were manually curated to break linkages at >30 cM, except in four cases where markers targeting alternate alleles of the same locus were within 40 cM in both parental maps. Rarely, markers mapping to a single scaffold were assigned to two different linkage groups. These discrepancies could be due to misassemblies in the draft somatic genome, genotyping errors or programmed genome rearrangements. A majority of these (totaling 2.8% of mapped scaffolds) were readily resolved by majority rule, with multiple mapped SNPs supporting an assignment to a specific linkage group. A smaller fraction of scaffolds could not be resolved by majority rule (1.5%) and these were placed arbitrarily. If misplaced, these scaffolds would be expected to contribute to (low level) background noise in the comparative maps, but when placed properly they should contribute to the signal of conserved synteny (again to a minor degree). Markers not incorporated into the map may represent distal portions of chromosomes, uninformative segregation phases or genotyping errors.

### Statistical analysis of **conserved** syntenic regions

Methods for identifying putatively orthologous loci are described elsewhere (Smith et al. 2013). Lamprey gene annotations used for this study were previously published and can be downloaded and browsed through the UCSC genome browser, http://genome.ucsc.edu/ (Kent et al. 2002; Smith et al. 2013). Operationally, a locus from the reference genome is considered to be orthologous to a lamprey gene if all three of the following conditions are met: 1) the t*blastn* (Altschul et al. 1990) alignment bitscore between the lamprey gene and a given locus from the reference genome is within 90% of the best alignment bitscore for that lamprey gene, 2) there are six or fewer paralogs detected in the lamprey genome and 3) there are six or fewer paralogs detected in the reference genome. This method circumvents many of the confounding effects of extreme substitution bias in the lamprey lineage, independent paralog loss between gnathostome and lamprey lineages, and the proximity of duplication and divergence events, all of which prevent accurate tree-based orthology assignment (Qiu et al. 2011; Mehta et al. 2013; Smith et al. 2013). In order to reflect the biological reality of whole genome duplication and subsequent evolutionary assortment of paralogs, “orthologs” refer to the set of loci that share a most recent common ancestor in the preduplicated/predivergence state. Counts of orthologs on all pairwise combinations of lamprey LGs and gnathostome (chicken: GCA_000002315.1 or human: GCA_000001405.1, http://www.ncbi.nlm.nih.gov) chromosomes were tabulated and compared to expected values based on random sampling from LGs and chromosomes with the same number of loci. Because these comparisons involve a large number of pairwise combinations of lamprey LGs and gnathostome chromosomes that often possess a small number of putative orthologs (i.e. most cells in the contingency table correspond to non-orthologous segments, especially in comparisons between species with extensive conservation of synteny), the distribution of orthologs was evaluated using a chi-square incorporating Yates’ correction for continuity and Bonferroni corrections for multiple testing.

Hypotheses as to mode of gene duplication were evaluated using goodness of fit tests, however, it should be noted that the small number of ancestral chromosomes presents a fundamental limitation to the power of these tests. Given that two rounds WGD are expected to yield ratios in excess of 1a:2d for chromosomes or chromosomal segments (expected values are zero for 1a:1d and 1a:2d) categorical test statistics become infinitely large for a test of goodness of fit to a model of 1a:4d (we use a conservative value of 1a:>2d for analyses performed here), therefore we applied a continuity correction to expected values in order to permit calculation of conservative G-tests (G_adj(c)_) (Sokal and Rohlf 1995). For models involving segmental duplication, expected distributions were based on Poisson distributions with mean values equivalent to the observed number of duplications divided by the number of ancestral chromosomes. Two sets of observed numbers were calculated, one based on the minimal set of duplications necessary to generate the observed patterns (i.e. 1a:4d is the product of two duplications) and another based on the maximal set of duplications necessary to generate the observed patterns (i.e. 1a:4d is the product of three duplications). Statistical tests of goodness of fit to models involving segmental duplication employed a G-test with Williams’ correction for small sample size (G_adj(W)_) (Sokal and Rohlf 1995).

## Data Access

Sequence data are deposited at the NCBI short read archives (http://www.ncbi.nlm.nih.gov/sra) under study number PRJNA232586.

## Acknowledgements

This work was funded by a grant from the National Institutes of Health (R01GM104123) and supported by startup funds from the University of Kentucky Department of Biology. We thank Marianne Bronner (California Institute of Technology) for providing access to lamprey husbandry facilities for production of embryos.

### Disclosure Declaration

The authors declare no competing interests.

